# Pre-trial exogenous visual flicker does not affect behavioral or EEG signatures of conflict processing

**DOI:** 10.1101/004747

**Authors:** Michael X. Cohen, Fraser William Steel

## Abstract

Activity in the theta frequency band (4-8 Hz) over medial prefrontal regions has been consistently implicated in top-down cognitive control processes, including recognizing and resolving response conflict. It remains an unanswered question whether these theta-band dynamics are a neural mechanism of cognitive control, or instead are epiphenomenal to the neural computational machinery but are useful indices of brain function. Here we addressed this question by attempting to boost conflict processing (or its EEG theta-band signatures) via pre-trial exogenous theta-band visual flicker. Although the flicker successfully entrained posterior brain networks, there were no effects of flicker on behavior or on EEG signatures of conflict processing. In this paper, we detail our attempts and discuss possible future directions for using exogenous flicker in the study of the role of endogenous brain oscillations in conflict processing.

---

Over the past ten years, evidence has amounted regarding the role of theta-band (4-8 Hz) oscillations in medial prefrontal regions and top-down cognitive control processes such as conflict resolution and error processing (Luu et al., 2004; Cohen, 2011; Cohen and Cavanagh, 2011; Cavanagh et al., 2012; Huster et al., 2013). This theta-band response seems to reflect an endogenous oscillatory process, because it is present even after removing the phase-locked event-related potential (ERP) (Cohen and Donner, 2013), is uncorrelated or weakly correlated with ERP measures of cognitive control such as the N2 and ERN (Cavanagh et al., 2012), and is a stronger predictor of conflict and trial-varying reaction times compared to ERP measures (Cohen and Donner, 2013).

Linking a cognitive process to an endogenous brain oscillatory dynamic facilitates a physiological interpretation of neurocognitive processes, because the neurobiological events that generate oscillations are fairly well understood (Wang, 2010), compared to the neurobiological events that underlie the ERP or the hemodynamic response measured by FMRI. Thus far, most studies investigating the role of theta activity in cognitive control have been correlative, meaning that brain activity is passively measured while human subjects perform cognitive control tasks. Correlative evidence is crucial for characterizing the relationship between theta oscillations and cognitive control. At this point, however, we believe that sufficient correlative evidence has amounted that warrants causal interventions to determine whether theta oscillations *per se* are a core mechanism of conflict processing, or whether theta oscillations are a useful index into cognitive control functioning but are not part of its neural computational machinery.

There are several ways to introduce causal interventions in human neuroscience research, including transcranial magnetic current stimulation (Thut and Miniussi, 2009; Thut et al., 2011b) or transcranial alternating current stimulation (Schutter and Hortensius, 2011). In this study, we tried to induce theta by exogenous visual flicker (Thut et al., 2011a). Exogenous flicker refers to flashing a visual (or any other sensory modality) stimulus at a specified rate in order to entrain brain rhythms. It is often used to “tag” the processing of specific stimuli (known as frequency tagging or the steady-state evoked potential), but there is some evidence that flicker may also be used to boost frequency band-specific processing (Thut et al., 2011a). This is an attractive approach for studying the role of brain oscillations in cognition, because it is simple, requires no invasive brain stimulation, and can be done on the behavioral level or combined with electromagnetic or hemodynamic brain imaging. On the other hand, it has been debated whether a flickering visual stimulus actually entrains neural oscillations or merely produces a series of phasic evoked potentials (Moratti et al., 2007; Capilla et al., 2011; de Graaf et al., 2013) that, due to the rhythmic nature of the visual stimulus in combination with the Fourier Theorem, would allow rhythmically timed non-oscillatory processes to be captured as an apparently oscillatory process.

Although visual stimulus flicker primarily entrains activity in visual cortex, it has been shown that flicker can also modulate activity in larger networks that extend beyond early visual cortex (Ding et al., 2006; Srinivasan et al., 2006). One example that is relevant to the present study is that theta-band flicker increases hemodynamic activity in medial frontal regions in addition to occipital regions (Srinivasan et al., 2007). Thus, the goal of the present study was to use a pre-trial visual flicker in the theta band to test whether that would enhance behavioral or EEG signatures of conflict processing. The control conditions included a random flicker and an alpha-band (∼10 Hz) flicker.

To preview the results, we found a very robust effect of the pre-trial flicker (including at the first and second harmonics), and a significant conflict effect, but no interactions between flicker rate and conflict processing on the behavioral or EEG levels. We also tried several post-hoc exploratory analyses in hopes of finding evidence in favor of our hypotheses; however we failed to identify any significant temporally lasting effects of the flicker on subsequent task processing. We are thus confident in, though disappointed by, this null finding. In the Discussion section, we outline some possible reasons for the null result, and suggest some future directions that might help optimize the use of exogenous visual flicker in the study of the role of prefrontal theta in cognitive control.

## Methods

### Participants

Twenty-one participants (16 female) with normal or corrected to normal vision were recruited through the University of Amsterdam psychology research participant system. Subjects signed informed consent before the experiment began, and received either course credit or money (14 Euros) for their participation. The experiment was approved by the University of Amsterdam psychology department ethics committee, and lasted approximately two hours, including preparation time and the task. Data were collected in winter/spring of 2010.

### The task

The task procedure is illustrated in Figure 1. We used a modified version of the Eriksen flanker task. The subjects’ primary task was indicate the central letter that was present in a string of five letters, as quickly as possible using a response box that is built into the chair. On each trial of this task, the target letter could be either the same (congruent) or a different (incongruent) from the flanking letter. Each block contained 200 trials based on letter-pairs X-Y, E-F, B-R, or M-N (thus, an incongruent trial would contain the letter string “X X Y X X”). Fifty percent of trials were incongruent. Response-to-target letter mapping was such that the earlier letter in the alphabet (X, E, B, M) was associated with a left thumb response, and the other letter with a right thumb response. This was done to avoid introducing conflict between the response hand and a spatial representation of the alphabet. Letters were printed in Arial font (size 24) with a single blank space between letters, and subjects sat with their eyes approximately 90 cm from the monitor (19 inches, 1024 × 768 resolution). A 1200 ms inter-trial-interval separated the button press on one trial, and the flicker onset of the next trial. Stimuli were on screen for 50 ms.

**Figure 1.**
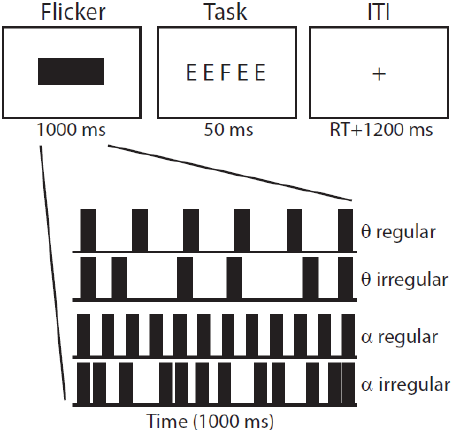
Task overview. The top row illustrates the visual stimuli on the screen and their durations (RT=reaction time). The bottom rows illustrate the timing of the four flicker conditions. The inter-flash-interval timing of the irregular flicker conditions was randomized on each trial, although there were an equal number of flashes in regular and irregular conditions.

### Visual flicker entrainment

Preceding each letter-pair on each trial, a solid black bar flickering at 6.7 Hz (theta) or 12.2 Hz (alpha) appeared at the center of the screen for 1000 ms (the background screen color was white). Subjects were instructed to fixate at the center of the screen before and during the flicker. They were also told that the flicker was task-irrelevant and was uncorrelated with the task conditions. The bar was 150 × 50 pixels (40 × 13 mm on our 19-inch monitor), and subtended 2.54 × 0.83 degrees visual angle. The monitor refresh rate was 60 Hz.

The control condition for the visual flicker was a flicker (same size as described in the previous paragraph) that had randomized inter-flash-intervals. Thus, the rhythmic nature of the flicker was abolished, although the number of flashes was the same as for the real flicker. Thus, we refer to “theta-regular” as the normal flicker, and “theta-irregular” as the randomized control-condition flicker. Flicker frequency was manipulated (pseudo-randomized) across blocks. During “regular” blocks, the stimulus onset was timed to be in-phase with the stream of flickers. Within each block, regular and irregular flicker trials were randomly interleaved, and present in 50% of trials, but each block would contain only theta flicker or only alpha flicker. Furthermore, there was only one letter-pair (X-Y, E-F, B-R, or M-N) per block. The relationship between letter-pair and flicker condition (theta vs. alpha) was randomized across subjects, but was constant within each subject. In practice, this meant that we constructed eight task blocks for all combinations of letter-pairs and flicker frequencies (e.g., X-Y theta and X-Y alpha), and each subject would perform four of these blocks (ensuring that each subject had all four letter-pairs once and two blocks of each flicker frequency).

Each of the four blocks began with practice trials so subjects could demonstrate that they understood the task instructions and the mappings between the letters and the correct response hand. Rest breaks were offered after every 50 trials, at which point subjects were presented with on-screen feedback regarding their behavioral performance (percentage correct and average reaction time, unrelated to the flicker condition and averaged over congruent and incongruent trials), and could continue by pressing a button.

### EEG and EMG data collection and analyses

Electrophysiological activity was measured from 64 electrodes placed according to the 10-20 system, acquired at a sampling rate of 512 Hz. Additionally, two horizontal electro-oculogram electrodes were used to record eye movements, and two electrodes were attached to the earlobes (used as a reference). All electrode sites were grounded through CMS/DRL electrodes (see www.biosemi.com for additional hardware details). Electromyographic (EMG) recordings were taken from the flexor pollicis brevis muscle of each thumb using a pair of surface electrodes, placed on a subject-by-subject basis approximately 5 mm apart on the thenar eminence.

Offline, data were high-pass filtered at 0.5 Hz and epoched between -1.5 and +3 seconds around each flicker onset. Trials containing muscle or other EEG artifacts were removed after visual inspection. Data from two subjects were excluded from analysis due to excessive EEG noise; thus, group-level analyses contained data from 19 subjects. Trials with eye-blinks occurring during the flicker period were rejected. Independent components analysis (ICA) was computing using the eeglab toolbox (Delorme and Makeig, 2004) for Matlab, and components clearly capturing blink artifacts or other non-brain-related artifacts were removed from the data prior to analyses. Error trials, post-error trials, and partial error trials were excluded from analyses. Partial errors are trials in which the subject twitched the muscle of the response hand corresponding to an error, when they pressed the correct button. These trials can contaminate “pure correct” trials, as we have detailed in our other recent studies (Cohen and Van Gaal, 2012; Cohen and van Gaal, 2013). EEG data were spatially filtered using the surface Laplacian; this minimized the possibility that flicker effects at electrode FCz reflected volume conduction from visual cortex (we often use the surface Laplacian and find that it helps to localize the conflict modulation of theta to midfrontal electrodes). For the analyses of conflict processing, we analyzed both the total power and the non-phase-locked power; the non-phase-locked power was obtained by subtracting the ERP from each trial before performing time-frequency decomposition. Our recent findings (Cohen and Donner, 2013) suggest that most of the conflict-related theta power is non-phase-locked. The total power was used for analyses on flicker effects.

Time-frequency decomposition was carried out using customized Matlab scripts. We applied complex Morlet wavelet convolution to obtain estimates of time-frequency-electrode power and phase. Details are presented in many of our other papers and in a textbook (Cohen, 2014). We extracted 20 frequencies between 2 Hz and 40 Hz. We focus on two measures: power (frequency band-specific squared amplitude; converted to decibels using a -300 to -100 ms pre-flicker baseline period) and inter-trial-phase-clustering (ITPC; the consistency of the timing of band-specific activity at each time-frequency point over trials).

### Statistics

Statistics on behavioral performance were performed using a three-way repeated-measures ANOVA on average reaction time (RT) or accuracy (separate ANOVAs for RT and accuracy). The factors were frequency (theta, alpha), flicker type (regular, irregular), and condition (congruent, incongruent). The ANOVAs were performed in SPSS software. For the RT analyses, only the RTs from correct trials were used.

Statistics on EEG data were performed in two ways. The first approach involved hypothesis-testing based on hypothesized results in defined a priori time-frequency-electrode windows. In this approach, data were tested as described above for the behavioral data from specified time-frequency windows that were motivated by results from orthogonal analyses (described where appropriate in the Results section).

The second approach for statistical evaluation of EEG data was to perform non-parametric permutation testing. This approach is useful when there are no strict hypotheses governing the time-frequency windows for analyses, and when correcting for multiple comparisons across all time-frequency points. At each of 1000 iterations, the effect of interest (EEG activity during incongruent compared to during congruent trials; or activity during regular compared to irregular flicker conditions) time-frequency map was multiplied by -1 for a random subset of subjects (this effectively randomizes A-B vs. B-A). Next, a t-statistic was calculated at each time-frequency point, the t-map was thresholded at p=0.05, and the cluster containing the largest sum of absolute magnitude t-values was stored; thus, after 1000 iterations, a distribution of maximum clusters expected under the null hypothesis could be created. In the final step of the analysis, the true t-statistic map was thresholded at p=0.05, and any supra-threshold clusters that were smaller than the 95% percentile of the null hypothesis cluster distribution were removed. This procedure corresponds to a p-value of 0.05, correcting for multiple comparisons across all time points and frequency bins. More details and justifications of this procedure can be found in Maris and Oostenveld (2007).

## Results

### Behavior

Behavioral results are illustrated in Figure 2. As expected, RTs were longer, and accuracy was lower, on incongruent compared to congruent trials. This was demonstrated by a significant main effect of trial condition (RTs: F_1,18_=21.26, p<0.001; accuracy: F_1,18_=20.46, p<0.001). There was also an unexpected but significant main effect of flicker frequency, such that subjects were significantly faster (F_1,18_=4.45, p=0.049) and more accurate (F_1,18_=6.02, p=0.025) in blocks with theta flicker compared to alpha flicker. However, there were no significant interactions between flicker condition and flicker regularity or conflict (all p’s>0.1). This main effect of flicker frequency is difficult to interpret because it was present in both the regular and irregular conditions; thus, having fewer flashes was associated with faster and more accurate responses, compared to having more flashes (alpha conditions).

**Figure 2.**
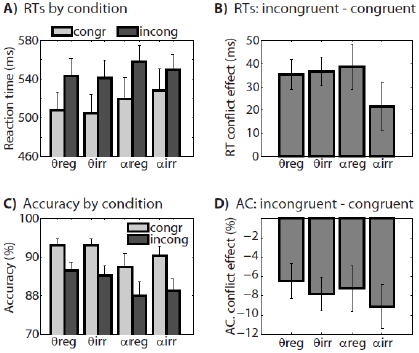
Behavioral results. Panels A and B illustrate reaction times (RTs), and panels C and D illustrate accuracy (AC). Panels A and C show results for each condition separately, and panels B and D show the conflict effect (incongruent minus congruent trials) separately for each flicker condition. Our hypothesized interaction between theta-band (θ) flicker and conflict processing was not supported by the data.

The hypothesized three-way interaction among flicker rate, flicker regularity, and conflict was not statistically significant for accuracy: F_1,18_=0.53, p=0.47. There was an unpredicted by significant three-way interaction for reaction times (F_1,18_=5.33, p=0.033), which was driven by a reduced conflict effect in the alpha-irregular condition. There was no effect for theta flicker.

### EEG flicker effects

We quantified the effect of flicker on the EEG signal as the difference in activity between theta regular and theta irregular, and between alpha regular and alpha irregular. Note that in this comparison, there is an equal number of congruent and incongruent trials in the subtraction, and that both regular and irregular flicker conditions contained an equal number of visual flickers (only the inter-flicker-timing differed). Although flicker effects are often shown in spectral power plots, our effects were quite robust, and we therefore show time-frequency plots to highlight the time course of the effects. Results are shown in Figure 3.

**Figure 3.**
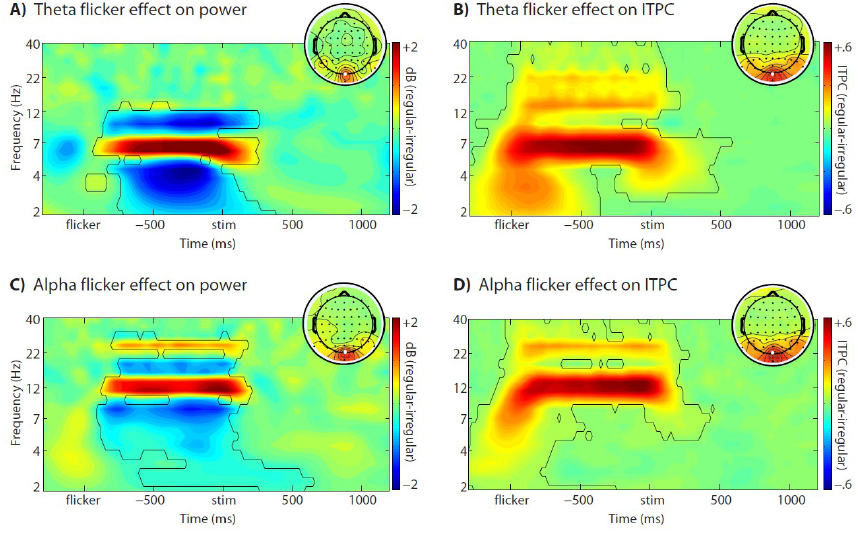
Effects of visual flicker (duration from “flicker” until “stim”) on EEG power and ITPC at electrode Oz (see white dot on topographical maps). All plots show the difference in activity between regular and irregular flicker conditions. Black contours outline regions of statistical significance (p<0.05, corrected for multiple comparisons over time-frequency points). Topographical map insets illustrate activity from -800 to 0 ms at 6.5 Hz or 12.2 Hz (color scaling is the same as for the time-frequency maps). The relative suppressions of power above and below the flicker frequencies are due to broadband increases of power during irregular flicker conditions.

It can be seen that both the theta and alpha flicker elicited increases in frequency band-specific activity. This was clear in both power and in ITPC. Increased power in several harmonics can also be seen in the ITPC plots. As expected for visual flicker, the effects were maximal at central occipital electrodes, with spatial peaks at electrodes Oz and Iz.

Because we had an a priori hypothesis concerning the effect of flicker over midfrontal regions, we also tested for significant changes in power and ITPC at electrode FCz. Because we did not want to restrict ourselves to testing for any specific time-frequency window, we tested all time-frequency pixels and applied a cluster-based correction for multiple comparisons (p<0.05; see Methods). This was done twice: Once for theta regular vs. theta irregular, and once for alpha regular vs. alpha irregular. We selected FCz for this analysis because many previous studies, and the present analyses (see next section), topographically localize response conflict effects to this electrode. There were no supra-threshold clusters for either theta-band or alpha-band flicker. Furthermore, visual inspection of the results did not suggest any nearly-significant results (data not shown).

### EEG conflict effects

In this analysis, we replicated a growing number of studies that show an enhancement of midfrontal theta-band power during high-conflict compared to low-conflict conditions. As expected, comparing all incongruent trials to all congruent trials (collapsing over all flicker conditions) revealed a selective enhancement of theta-band power that was localized to electrode FCz (Figure 4). This finding demonstrates that the typical conflict effect is observed, despite the slightly unusual design involving the pre-trial flicker.

**Figure 4.**
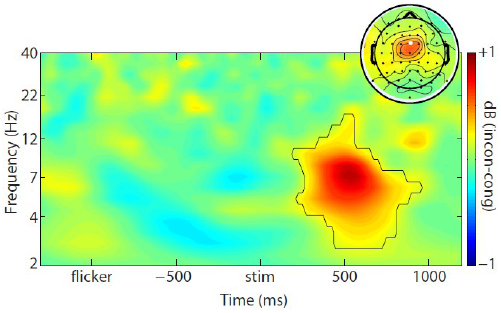
Conflict effect on EEG time-frequency power at electrode FCz (see white electrode in topographical map inset). The black contour outlines a region that exceeded statistical significance at p<0.05, corrected for multiple comparisons over all time-frequency points at this electrode. Topographical map illustrates the conflict effect from 400-800 ms and 4-8 Hz. The color scaling is the same for both the time-frequency power plot and the topographical map.

### EEG conflict effects modulated by flicker frequency

In this set of analyses, we examined whether EEG signatures of the conflict effect were modulated by the pre-trial flicker. In all analyses, results were compared between theta regular vs. theta irregular, and between alpha regular vs. alpha irregular.

We first performed a region-of-interest analysis, in which the conflict effect on theta power was examined separately according to the pre-trial flicker condition (note that the time-frequency window was selected based on the conflict effect, averaging over all pre-trial flicker conditions). No hypothesized interaction effects emerged (all p’s>0.2) (Figure 5a-b). Next, we performed an exploratory analysis in which we tested the influence of the theta flicker on conflict-related activity at all time-frequency points at electrode FCz. Again, no significant effects emerged (Figure 5c).

**Figure 5.**
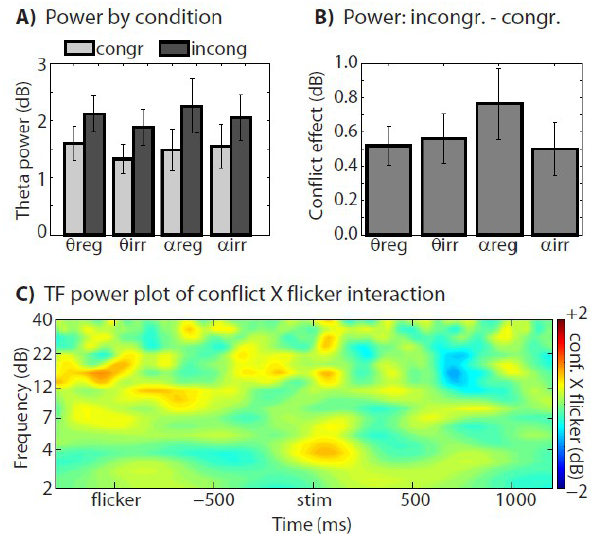
This figure shows the null results of the main hypothesis of the study. We predicted that the pre-trial theta-band flicker will enhance EEG signatures of cognitive control. This was not observed, either in the time-frequency window selected based on the main effect of conflict (see Figure 4) shown in panels A and B, or in an exploratory analysis over all time-frequency points at electrode FCz (panel C).

### Additional post-hoc analyses to search for flicker-modulated conflict effect

We tried several additional post-hoc analyses in search of an effect of theta flicker on conflict-related EEG signatures. None of these analyses revealed any convincing effects in our hypothesized directions, but we describe the methods here briefly to inform the reader of what additional analyses were conducted. Results are not shown because there were no significant or visually compelling results in the predicted directions.

First, we performed a principle components analysis on the stimulation period data, extracted the first two principle components, and applied a time-frequency analysis as for the electrode-level data. Although the first two components fairly cleanly dissociated the pre-trial flicker effects and the task-related “cognitive” effects, there were no significant (or even visually compelling) effects of flicker condition on conflict-related time-frequency power.

Second, we performed single-trial brain-brain correlation analyses. The idea here was to quantify the effect of flicker on each trial, and use this in a single-trial correlation analysis. The effect of single-trial flicker was estimated as the amplitude of the Fourier coefficient corresponding to that trial's flicker frequency (theta or alpha) on activity at Oz. That vector over trials was then correlated with cross-trial time-frequency power dynamics. Although we observed large correlation coefficients with power at the flicker frequency during the flicker period, there were no significant correlations during the stimulus/response period, and there were no differences between congruent and incongruent trials.

Third, we computed phase synchronization between FCz and Oz, and between FCz and F6. This analysis was motivated by our previous studies that highlight the role inter-regional theta-band phase synchronization in conflict processing and error monitoring (Cavanagh et al., 2009; Nigbur et al., 2011). Although conflict modulations of phase synchronization were in expected directions (enhanced connectivity for incongruent compared to congruent trials), there were no modulations by flicker condition.

Fourth, we applied Granger-based directed connectivity analyses to examine whether “bottom-up” (Oz-->FCz) or “top-down” (FCz-->Oz) conflict-related connectivity was modulated by flicker. Although we observed increased top-down conflict-related connectivity that mirrored our previous findings on post-error-related modulations of directed connectivity (Cohen et al., 2009), there were no compelling modulations of this effect by the flicker condition.

Fifth, we computed correlations between the flicker effect on RTs and the flicker effect on the EEG. Here the idea was that if the flicker successfully modulated conflict processing to different degrees in different subjects, correlation analyses might reveal this relationship (this has been demonstrated for TMS effects on posterior alpha-band oscillations; Hamidi et al., 2009). Although the correlation coefficient was positive, it was not statistically significant (r=0.18, p=0.46).

In theory, there are many many other possible ways to analyze EEG data that we could have tried, but given the consistent null results in the hypothesis-driven analyses and in the exploratory analyses, we decided that further analyses would be unlikely to yield any effects. Furthermore, if we were to obtain a significant effect in one out of, say, ten analytic approaches, that result would be difficult to believe.

## Discussion

Because the statistically significant effects that we report are mainly replications, we will keep our discussion here focused on the null results of the main hypotheses. Readers interested in how large-scale brain networks may become entrained by stimulus flicker may start with the following references (Srinivasan et al., 2006; Di Russo et al., 2007; Vialatte et al., 2010), and readers interested in the role of midfrontal theta in cognitive control may find the following references a good starting point (Cohen, 2011; Cavanagh et al., 2012; Cohen and Van Gaal, 2012; Cohen and van Gaal, 2013).

There are several possible reasons that might explain our null result. Of course, it is possible that theta-band oscillations play no causal role in cognitive control, but we prefer to tentatively maintain our initial hypothesis and instead focus on methodological limitations of the present study and suggestions for follow-up research.

One reason for the null result may be that the flicker occurred prior to the trial onset, rather than during the task. The extent to which brain oscillations can remain entrained after the flicker ends remains unclear. A review paper suggested that the effect of flicker on brain oscillations lasts only as long as the flicker remains on-screen (Thut et al., 2011a). On the other hand, lingering entrainment effects have been reported to last as long as 1.6 seconds (Sakamoto et al., 1993) and with consequences for behavior several cycles after flicker offset (Spaak et al., 2014). Having a flicker during stimulus presentation in the Flankers task (and other typical cognitive control tasks) presents two difficulties. First, RTs tend to be fast in these kinds of tasks, which means that there may be only two to four theta-band flashes during stimulus presentation. This is likely to be too few flashes to entrain brain networks. This is not to say that flicker cannot be used during cognitive control tasks; higher-frequency flicker has been used for frequency tagging to isolate processing of targets and distracters (Scherbaum et al., 2011; Gulbinaite et al., 2014). Second, because RTs are variable, the number of flashes will vary across trials, and this will furthermore correlate with conflict processing because more conflict processing generally means longer RTs (and thus, more flashes). These were the reasons why we decided to try a pre-stimulus flicker.

Thus, an idea for a follow-up study could be that the stimuli or background flicker both before and during stimulus presentation. This may promote tonic rather than phasic oscillatory activity, although it has been shown that resting state theta power recorded prior to a block of trials (this might reflect a tonic theta state) predicts task-related ERPs indices of conflict, at least for internally guided conflict such as preference judgments (Nakao et al., 2012).

A second possible reason for the null result is that the flicker was task-irrelevant. Indeed, subjects were instructed that the flicker provided no information regarding the upcoming trial. We designed the task this way to test for an effect of band-specific entrainment without any potential confound of flicker-rate-modulations of task preparation or cognitive set. Thus, another idea for a follow-up study would be to have the flickering stimulus be task-relevant (and perhaps even cognitive control-relevant). It is possible that a task-relevant flicker would be more deeply processed, and thus might entrain more wide-spread brain networks. A task-related flicker could involve the subject performing a secondary task on the flicker, for example by reporting trials that contain “oddball” fast flickers. Indeed, it did not appear from the present results that the flicker was processed more anteriorly than visual cortex. This contrasts with previous studies that showed parietal and frontal entrainment by visual flicker (Srinivasan et al., 2006, 2007). We speculate that the main difference is the size of the stimulus—our stimuli were fairly small while previous studies used large (in some cases, “ganz-field” or whole-field) high-contrast stimuli. It is possible that larger stimuli entrain networks beyond visual cortex.

Yet another possibility is to enhance the entrainment by using a multimodal flicker, such as simultaneous visual and auditory flickers. Because the computations in medial frontal cortex are thought to be fairly high-level and not specifically linked to any particular input or output modality (Ridderinkhof et al., 2004; Ridderinkhof et al., 2011), it is possible that a multimodal flicker will facilitate the entrainment of larger-scale brain networks, possibly including the medial frontal cortex.

A third possible reason for the null result is that the stimulation frequencies were not custom-tailored to each individual subject. Intracranial empirical and modeling studies have suggested that rhythmic brain stimulation is optimal when there is a match between the exogenously applied rhythm and the endogenous brain oscillation (Reato et al., 2013) (these studies used electrical stimulation, but the same principal may apply to visual stimulus flicker). In other words, 6 Hz stimulation might have little effect for an individual who has a 7.5 Hz endogenous theta.

Despite the null findings, we would like to conclude on a positive note. There are many methods to study the causal roles of oscillations in human brain function, and there is tentative evidence from other cognitive domains that sensory flicker may be a successful method for experimenter-controlled causal modulations of brain oscillations (Thut et al., 2011a). Testing for causal roles of brain oscillations in cognition is one of the important challenges in cognitive electrophysiology over the next decade. We are optimistic that stimulus flicker can be used to provide evidence for a causal role of theta-band oscillations in cognitive control process, and we hope that we and others will learn from the present null effects.

## Acknowledgements

This research was funded by a VIDI grant awarded to MXC from the Netherlands Organization for Scientific Research (NWO).

